# Association of *Bacteroides acidifaciens* relative abundance with high-fibre diet-associated radiosensitisation

**DOI:** 10.1101/846436

**Authors:** Chee Kin Then, Salome Paillas, Xuedan Wang, Alix Hampson, Anne E Kiltie

## Abstract

**Background:** Patients with pelvic malignancies often receive radiosensitising chemotherapy with radiotherapy to improve survival, however this is at the expense of increased normal tissue toxicity, particularly in elderly patients. Here we explore if an alternative, low-cost and non-toxic approach can achieve radiosensitisation in mice transplanted with human bladder cancer cells. Other investigators have shown slower growth of transplanted tumours in mice fed high-fibre diets. We hypothesised that mice fed a high-fibre diet would have improved tumour control following ionising radiation (IR) and that this would be mediated through the gut microbiota.

**Results:** We investigated the effects of four different diets (low fibre, soluble high fibre, insoluble high fibre and mixed soluble/insoluble high fibre diets) on tumour growth in immunodeficient mice implanted with human bladder cancer flank xenografts and treated with ionising radiation, simultaneously investigating the composition of their gut microbiomes by 16S rRNA sequencing. A significantly higher relative abundance of *Bacteroides acidifaciens* was seen in the gut (faecal) microbiome of the soluble high fibre group, and the soluble high fibre diet resulted in delayed tumour growth after irradiation compared to the other groups. Within the soluble high fibre group, responders to irradiation had significantly higher abundance of *B. acidifaciens* than non-responders. When all mice fed with different diets were pooled, an association was found between the survival time of mice and relative abundance of *B. acidifaciens*. The gut microbiome in responders was predicted to be enriched for carbohydrate metabolism pathways and *in vitro* experiments on the transplanted human bladder cancer cell line suggested a role for microbial-generated short-chain fatty acids and/or other metabolites in the enhanced radiosensitivity of the tumour cells.

**Conclusions:** Soluble high fibre diets sensitised tumour xenografts to irradiation and this phenotype was associated with modification of the microbiome and positively correlated with *B. acidifaciens* abundance. Our findings might be exploitable for improving radiotherapy response in human patients.

## Background

Patients with pelvic tumours, including bladder, cervix and rectal cancers, who are receiving radiotherapy are often given additional radiosensitising chemotherapy to improve cure rates, at the expense of increased toxicity in local organs and tissues (1, 2). With an ageing population, new approaches to radiosensitisation are urgently required. One such approach might be to modify the intake of dietary fibre by supplements before and during radiotherapy or current standard chemoradiation schedules, which would be a very cost-effective strategy, not expected to add to normal tissue toxicity (3, 4).

Wei *et al* showed slower growth rates of subcutaneous lymphoma xenografts in mice fed a high fibre diet (8%) compared to mice on a low-fibre diet, with similar findings in both immune-deficient and immune-competent models (5). This was associated with increased plasma and tumour butyrate levels, but the authors did not investigate the effects of the diet on the gut microbiome.

Dietary fibre manipulation can very rapidly alter the human gut microbiome, with changes in faecal short chain fatty acid (SCFA) levels seen only one day after the diet reaches the distal gut (6). Dietary fibre can also mediate systemic immune effects (7), as can the microbiota-derived SCFAs (8, 9). Furthermore, dietary fibre structures align with phenotypes of specific microbes that differ in their metabolic pathways (10). SCFAs are known to confer anti-cancer effects (5, 8, 9). Other metabolites, including small intermediate and end by-products of endogenous metabolic pathways, products of microbe-host co-metabolism and exogenous signals arising from diet, drugs and other environmental stimuli might also be important (11).

We hypothesised that systemic effects of altered metabolites secreted by gut bacteria on tumours, due to dietary fibre modification, could be exploited in conjunction with ionising radiation (IR) to achieve radiosensitisation.

The aims of this study were to examine the impact of the diet on the microbiome before and after irradiation, and to correlate diet-induced microbiome changes with tumour growth and response to radiation treatment.

## Results

### The environmental microbiome had minimal impact on gut microbiome analysis

Female CD1 nude mice were injected subcutaneously with RT112 bladder carcinoma cells, and, at the same time, they commenced a modified diet, namely, one of the following: low dietary fibre (0.2% cellulose), low fibre with butyrate in drinking water, high soluble fibre (10% inulin), high insoluble fibre (10% cellulose) and high mixed fibre (5% cellulose, 5% inulin) (Figure 1A and Additional file 1: Table S1). We quantified bacterial loads using PCR and gel electrophoresis, compared to known numbers of *E. coli* colony forming units (CFU). Our mouse samples contained more than 10^4^ bacterial CFUs which appeared to override contaminating species in the sample microbial communities (Figure 1B). The PBS negative control was processed from the start of the DNA extraction identically to the luminal contents and tissue samples. The amount of nucleic acid detected in the PBS negative controls was extremely low, compared to that in the gut microbiota (Figure 1C). Furthermore, the community microbiome in this negative control differed markedly from the gut microbiome of the mice (Figure 1D). Therefore, the environmental microbiome had minimal impact on the analysis of the gut microbiomes of interest in this study.

**Figure 1.**
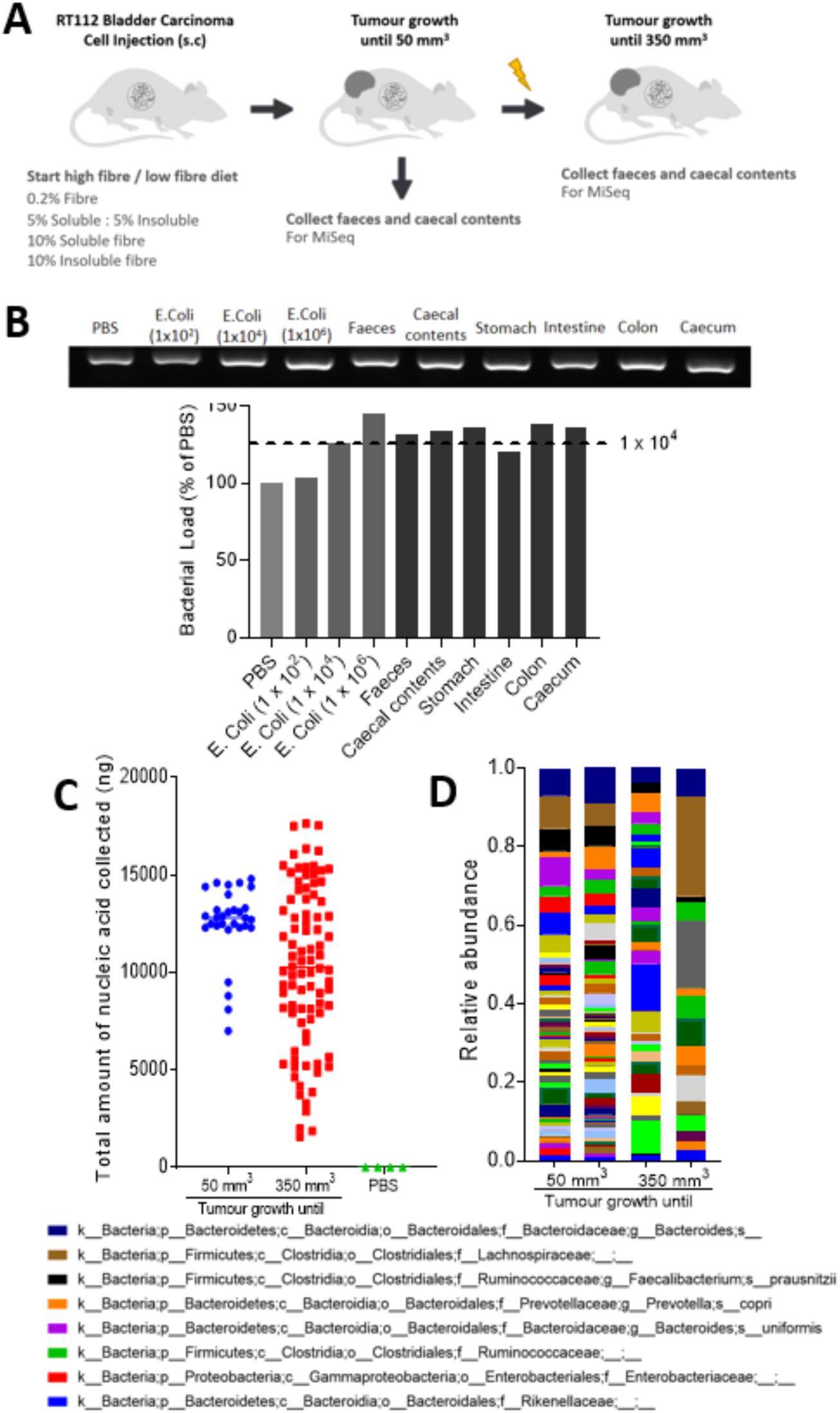
The environmental microbiome had minimal impact on gut microbiome analysis. (A) Two microbiomes were analysed from the intestinal tract, namely, faecal and caecal content samples collected when tumours reached 50 mm^3^ and 350 mm^3^ respectively. (B) Quantification of bacterial load from different tissue and luminal contents from mice, with *E. Coli* (1 × 10^2^, 1 × 10^4^, 1 x10^6^ CFUs) as controls (n=1 mouse). (C) Comparison of the total amount of nucleic acid quantified by PicoGreen assay in all samples collected when the tumours reached 50 mm^3^ and 350 mm^3^. (D) Common bacterial taxa at the species level in 4 samples of PBS, as negative controls of DNA extraction by 16S rRNA sequencing.

### The landscape, diversity and enrichment of bacterial taxa in the gut microbiome samples collected when the tumours reached 50 mm^3^

In samples collected when the tumours reached 50 mm^3^ and 350 mm^3^, the faecal (hereinafter referred to as “gut microbiome”) and caecal microbiome were found to have similar bacterial components (Additional file 1: Figure S1). In mice culled when their tumours reached 50 mm^3^, butyrate levels in the faeces, measured by high-performance liquid chromatography (HPLC), were found to be in the mM range, and were generally higher in the low fibre with butyrate and high soluble fibre groups (p=NS; Additional file 1: Figure S2A). The mean time for tumours to reach 50 mm^3^ was 12.8 ± 1.4 days (Additional file 1: Figure S2B).

In faeces collected from culled mice when the tumours reached 50 mm^3^, abundance analysis revealed the five bacterial taxa with the highest abundance were *Bacteroides acidifaciens, Parabacteroides, Akkermansia muciniphila, Lachnospiraceae* and *S24-7* (Figure 2A). In terms of alpha diversity, the soluble high fibre group had a lower Shannon’s index (p<0.001) (Figure 2B). This could be due to the higher abundance of *B. acidifaciens*, which lowered the diversity within groups. In terms of beta diversity, principal coordinate analysis showed a notable cluster effect among different groups, which indicates that samples within groups were more similar to each other than to those from the other groups (Figure 2C). This suggested that the gut microbiome was indeed modified in this study, which might be a diet-effect or a cage-effect (see later). Regarding the abundance of specific taxa in different diet groups (Figure 2D), the high soluble fibre diet significantly increased *B. acidifaciens* abundance (p<0.001); the low fibre diet increased *Parabacteroides* abundance (p<0.001), the low fibre diet with added butyrate increased *Akkermansia muciniphila* abundance (p<0.001), and the high mixed fibre diet increased *Lachnospiraceae* abundance (p=0.005).

**Figure 2.**
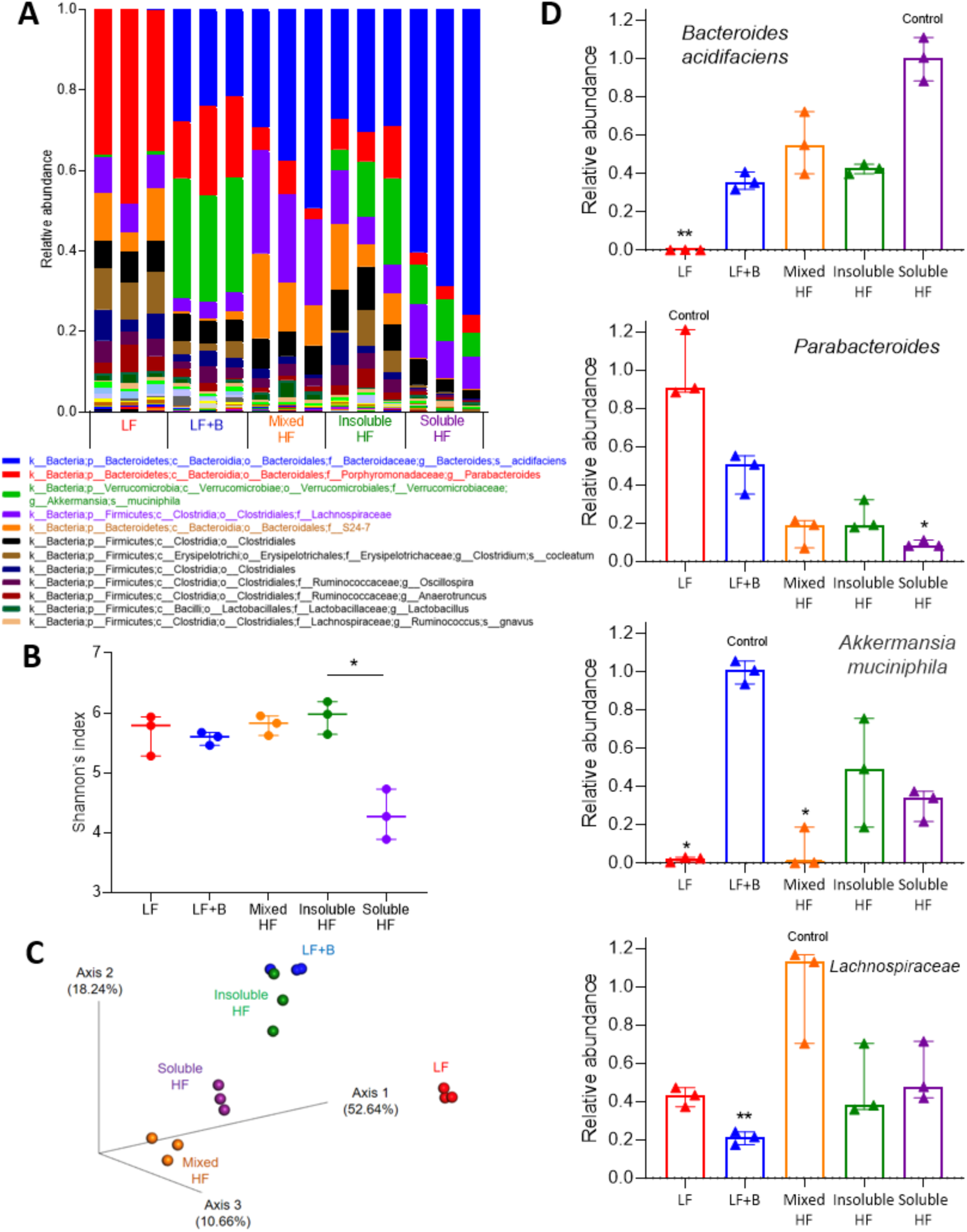
Dietary fibre shapes the baseline gut microbiome when tumours reached 50 mm^3^. (A) Stacked bar plot of phylogenetic composition of common bacterial taxa at the species level when tumours reached 50 mm^3^. Faeces were collected from mice fed with low fibre, low fibre with butyrate, high mixed fibre, high insoluble fibre and high soluble fibre diets (n=3 for each group). (B) Shannon’s index of gut microbiomes by Kruskal-Wallis test. Error bars represent the interquartile range of diversity scores. (C) Principal coordinate analysis of gut microbiomes using Bray-Curtis dissimilarity. A notable clustering effect by diet was seen in the gut microbiome. (D) Differentially abundant taxa when the tumours reached 50 mm^3^. All comparisons among different diet groups was performed by Kruskal-Wallis test and Dunn’s multiple comparison tests. All tests compared each median with the ‘control’ denoted. The diet with the highest abundance of a taxa was denoted as the control. *P<0.05; **P<0.01; ***P<0.001.

### The landscape, diversity and enrichment of bacterial taxa in the gut microbiome samples collected when the tumours reached 350 mm^3^

When the tumours reached 350 mm^3^, abundance analysis of the gut microbiome of both IR and non-IR cohorts revealed that the top 6 bacterial taxa with the highest abundance were *S24-7, Akkermansia muciniphila, Bacteroides, Lachnospiraceae, Clostridiales* and *B. acidifaciens* (Figure 3A). In terms of alpha diversity, the soluble HF group had a significantly lower Shannon’s index (p<0.001 for Kruskal-Wallis test) (Figure 3B). In terms of beta diversity, principal coordinate analysis using Bray-Curtis dissimilarity showed a notable clustering effect among different groups, which indicates that samples within groups were more similar to each other than to those from the other groups (Figure 3C). The composition of the gut microbiome continued to evolve on the diets to the time the tumours reached 350 mm^3^, regardless of whether the mice were irradiated or not. The taxonomic cladogram of LEfSe (Linear discriminant analysis Effect size) of the gut microbiome showed that the high soluble fibre diet increased relative abundance of *S24-7* (Additional file 1: Figure S3). *B. acidifaciens*, an acetate-producing bacteria (12, 13), was found to be of highest abundance in all but the LF diet group at 50 mm^3^, but by 350 mm^3^ was evenly distributed across the different diet groups, except the soluble HF group treated with radiation (p=0.200 for Kruskal-Wallis test) (Figure 3D).

**Figure 3.**
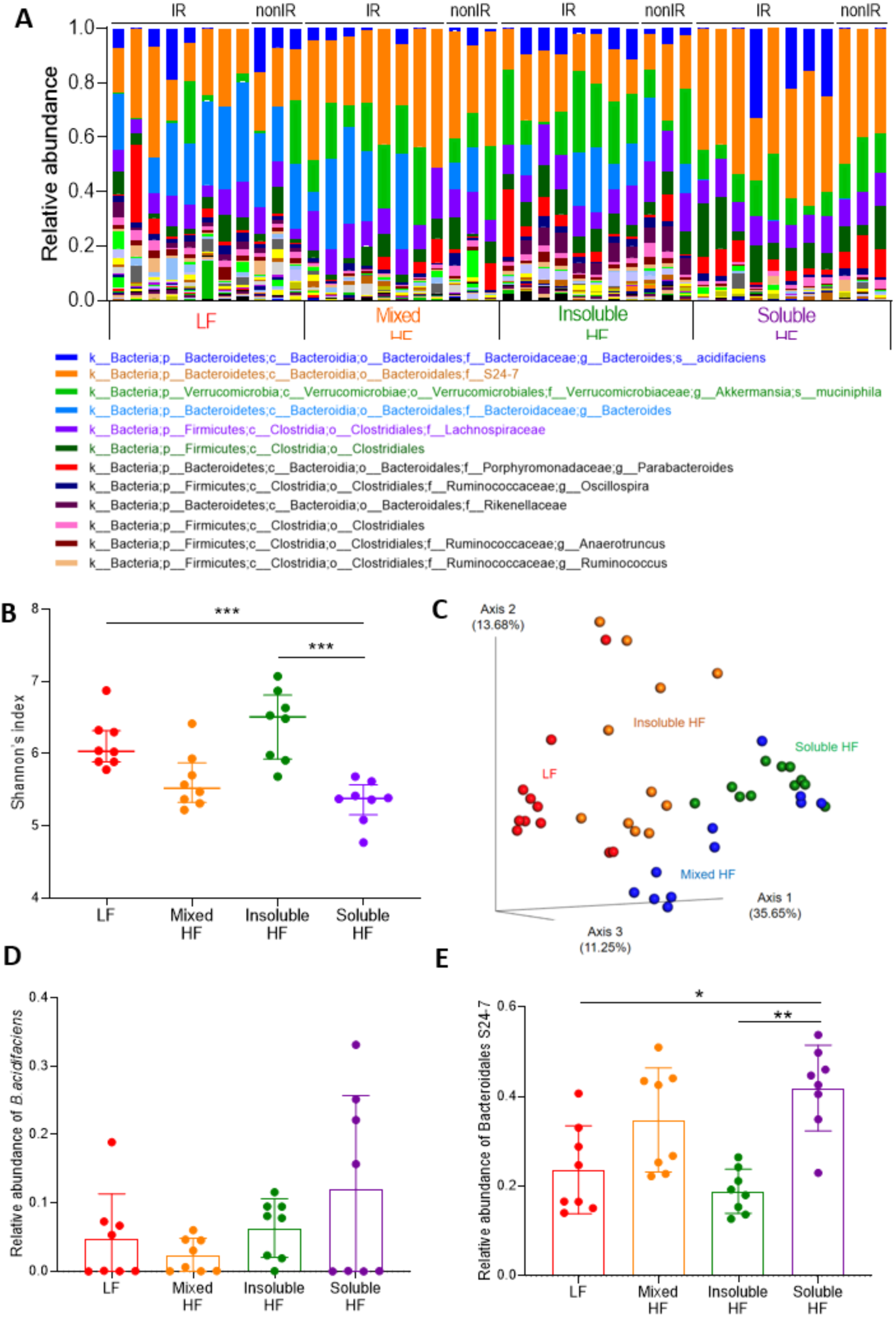
Composition of the gut microbiome when tumours reached 350 mm^3^. (A) Stacked bar plot of the phylogenetic composition of common bacterial taxa at the species level when tumours reached 350 mm^3^. Samples were collected from mice fed with low fibre, high mixed fibre, high insoluble fibre and high soluble fibre diets (n=8 for each group).(B to E) show results for irradiated mice only. (B) Shannon’s index of gut microbiomes by Kruskal-Wallis test. Error bars represent the interquartile range of diversity scores. (C) Principal coordinate analysis of gut microbiomes using Bray-Curtis dissimilarity. Relative abundance of (D) *B. acidifaciens* and (E) *Bacteroidales S24-7* in mice with or without irradiation. *P<0.05; **P<0.01; ***P<0.001.

*Bacteroidales S24-7* was the highest abundance bacterial taxa in the second cohort. Its relative abundance was significantly higher in the mixed HF and soluble HF groups compared to the LF and insoluble HF group (p=0.001 for Kruskal-Wallis test) (Figure 3E).

### Soluble high fibre causes increased growth delay in irradiated bladder cancer cell xenografts

To investigate the effect of different diets on the tumour response in mice irradiated when the tumour had grown to 50 mm^3^, tumour growth was monitored to 350 mm^3^. Slopes of the tumour growth curves were obtained using linear regression to indicate the tumour progression rates (Figure 4A). The high soluble fibre diet group had the slowest tumour growth rate. The slopes were 4.4 ± 1.3 for LF, 16.1 ± 1.7 for mixed HF, 28.7 ± 1.3 for insoluble HF and 0.4 ± 1.5 for soluble HF (Figure 4A and Additional file 1: Figure S4 for individual irradiated mouse tumour growth curves). Kaplan Meier survival curves for time to treble tumour volume showed that the soluble HF group had the longest median survival time (7.5 days for LF, 7 days for mixed HF, 10 days for insoluble HF, 11.5 days for soluble HF; p=0.005, log-rank test) (Figure 4B).

**Figure 4.**
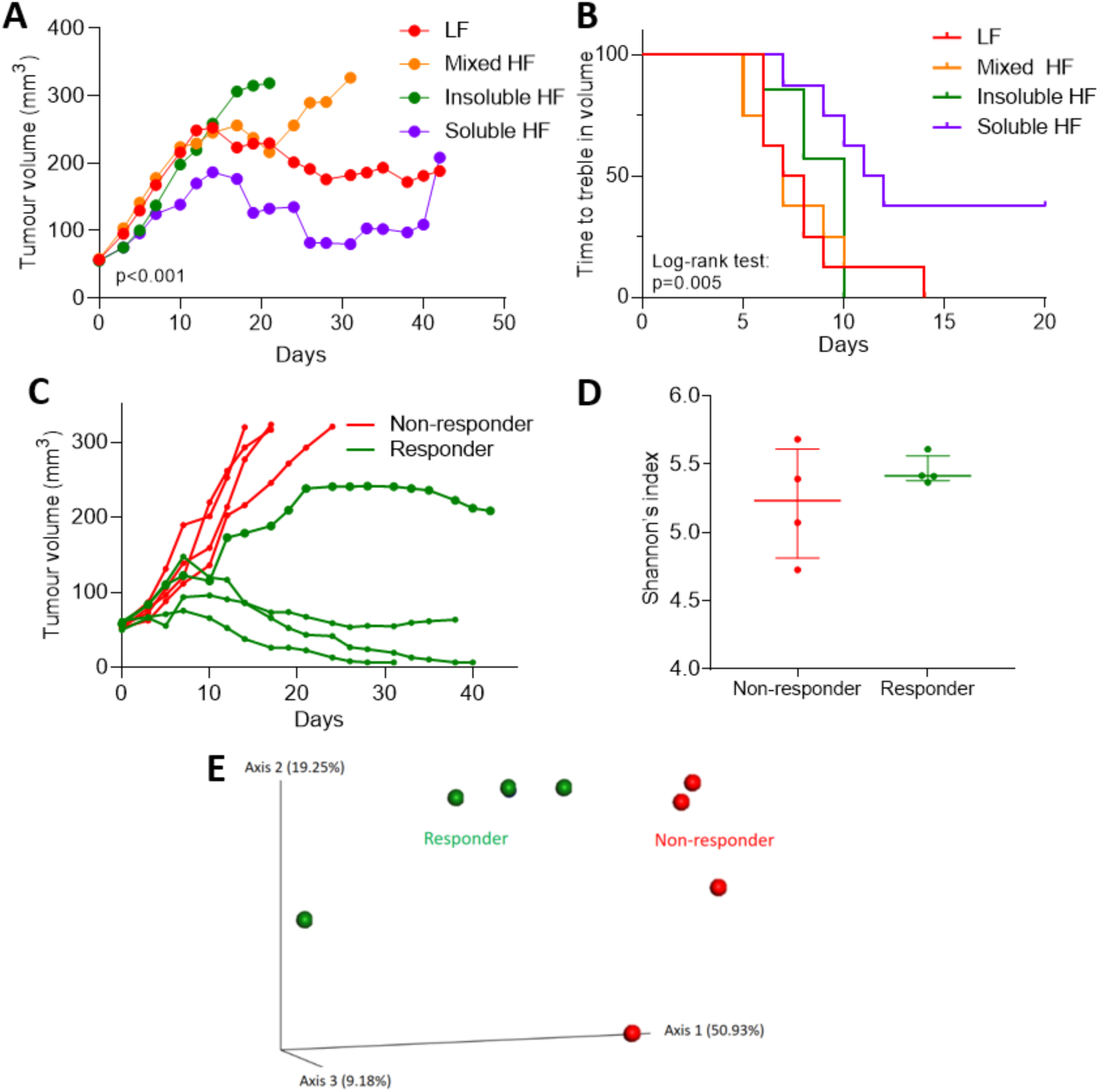
Soluble high fibre causes increased growth delay in irradiated bladder cancer cell xenografts and responses are influenced by gut microbiota composition. (A) Tumour growth in RT112 flank xenografts irradiated with 6 Gy IR, in mice fed low fibre, high mixed fibre, high insoluble fibre and high soluble fibre diets (n=8 for each group). Tumour curve slopes were calculated by linear regression to represent tumour growth rates and compared by ANOVA. (B) Kaplan–Meier survival curves for mice showing plots of time to treble tumour volume. (C) Mice in the soluble HF group were stratified into responders and non-responders based on tumour radiation response. (D) Shannon’s index of gut microbiota in responders and non-responders by Kruskal-Wallis test. Error bars represent the interquartile range of diversity scores. (E) Principal coordinate analysis of gut samples (n=8) in the soluble HF group by response using Bray-Curtis dissimilarity.

Among the eight mice fed the soluble high fibre diet, four mice were classified as responders as, using linear regression, they had shallower slopes to the tumour growth curves, namely, 7.3 ± 1.2, −0.9 ± 0.6, −4.8 ± 0.5, −5.2 ± 0.9. The other four mice were classified as non-responders with steeper slopes to the tumour growth curves, namely 34.6 ± 3.0, 31.1 ± 2.6, 23.3 ± 1.0, 33.8 ± 2.9 (Figure 4C). In terms of alpha diversity, there was no significant difference in Shannon’s index between responders and non-responders (Figure 4D). In terms of beta diversity, principal coordinate analysis of Bray-Curtis dissimilarity showed the gut microbiome of responders and non-responders were more similar within groups than between groups (Figure 4E).

### Differences in composition of the gut microbiome between responders and non-responders

Linear discriminant analysis showed that mice responding to the soluble high fibre diet with a slower tumour growth rate were enriched with *Bacteroidaceae (f), Flavobacterium (g), Flavobacteriales (o), Lactococcus (g), Streptococcus (g), Streptococcaceae (f), Allobaculum (g), Erysipelotrichales (o)*. The non-responding tumour-bearing mice were enriched with *Bifidobacterium (g), Bidifobacteriaceae (f), Bifidobacteriales (o), Parabacteroides (g), Porphyromonadaceae (f), Lactobacillus (g), Lactobacillaceae (f)*, and *Lactobacillales* (Figure 5A). In terms of effect size, *B. acidifaciens (sp)* and *Bacteroidaceae (f)*, had the largest enrichment in responders, and *Parabacteroides (g)*, and *Porphyromonadaceae (f)*, had the largest enrichment in non-responders (Figure 5B). To further explore these findings, the discrete false-discovery rates within all taxonomic levels were calculated (Figure 5C). In responders, *B. acidifaciens* species and the *Allobaculum* genus and in non-responders *Lactobacillus* and *Parabacteroides* genera had p-values <0.05 The *B. acidifaciens* abundance was significantly higher in responders than that in non-responders (p=0.029 (Figure 5D), while the *Bacteroidales S24-7* abundance was similar between responders and non-responders in the soluble HF group (p=0.200) (Figure 5E).

**Figure 5.**
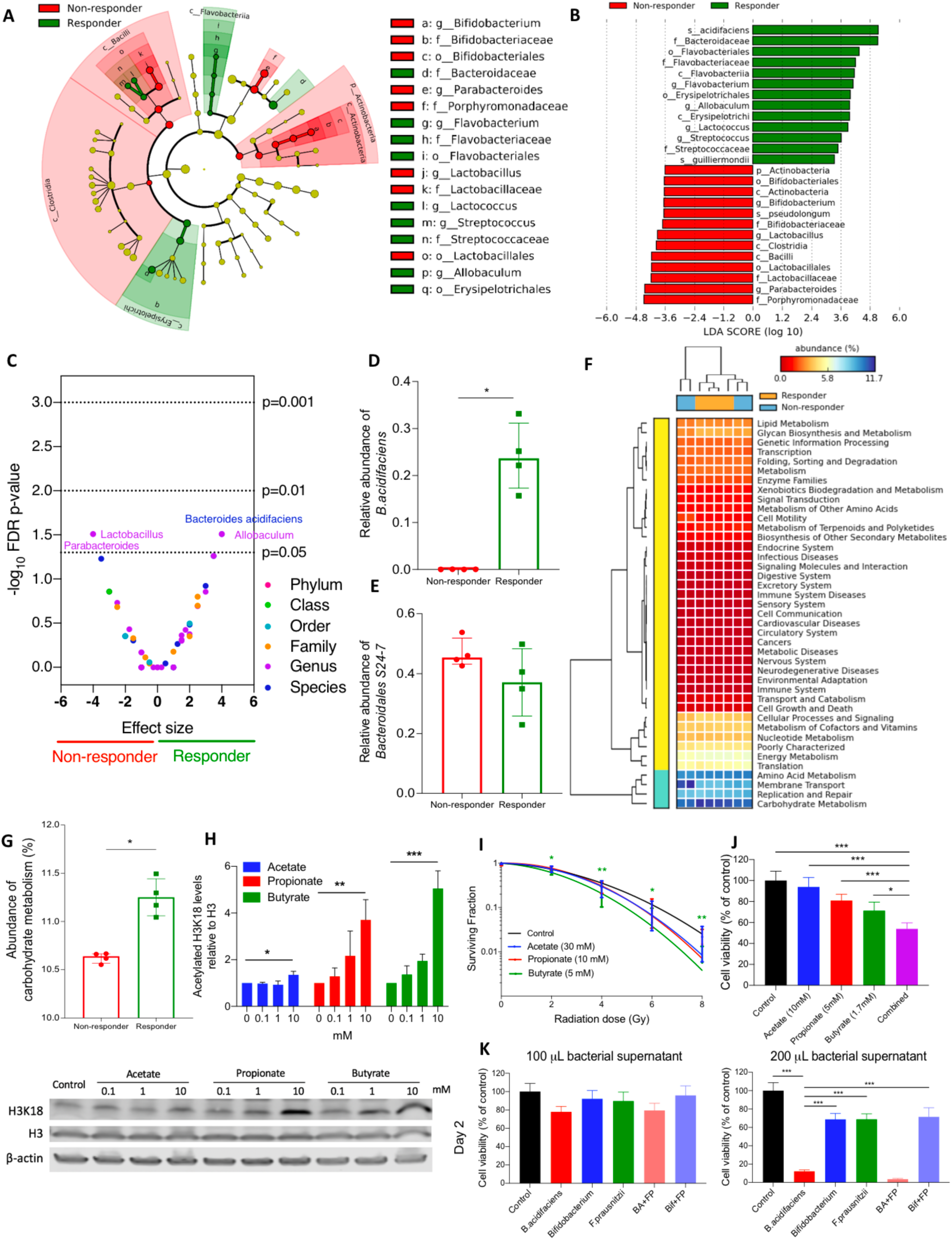
Differences in composition of the gut microbiome between responders and non-responders. (A) Taxonomic cladogram from LEfSe showing differences among taxa between responders and non-responders in the soluble HF group. Dot size is proportional to the abundance of the taxon. (B) Linear discriminant analysis (LDA) scores computed for differentially abundant taxa in the microbiomes of responders (green) and non-responders (red). Length indicates the effect size associated with a taxon, p=0.05 for Kruskal-Wallis test. (C) Discrete false-discovery rate of different abundant taxa in responders and non-responders in the soluble HF group. Differential abundance within all taxonomic levels in responders versus non-responders by Mann-Whitney U test. Dots are overlapping between *Bacteroides acidifaciens* and *Allobaculum*, and between *Lactobacillus* and *Parabacteroides*. Relative abundance of (D) *B. acidifaciens* and (E) *Bacteroidales S24-7* and in responders and non-responders in the soluble HF group. (F and G) Metagenomic functional prediction by PICRUSt of the gut microbiome in responders (n=4) and non-responders (n=4) in the soluble HF group with reference to the KEGG database level 2. Columns represent mice (responders, orange; non-responders, blue), and rows represent enrichment of predicted KEGG pathways (red, low enrichment; yellow, medium enrichment; blue, high enrichment). (H) Western blot analysis of histone acetylation levels of RT112 cells treated with SCFAs (N=3). (I) Linear quadratic survival curves of IC10-treated RT112 cells with receiving irradiation of 0-8 Gy (N=3). (J) Cell survival analysis of RT112 cells treated with single SCFA and combined SCFAs mixture (N=3). Combined (purple bar) denotes SCFA mixture of 10 mM acetate, 5 mM propionate and 1.7 mM butyrate. (K) Reduced cell survival of RT112 cells by bacterial supernatants at day 2 (N=1). *BA+FP* denotes the cross-feeding of *B. acidifaciens* and *F. prausnitzii*, while *Bif+FP* denotes the cross-feeding of *Bifidobacterium* and *F. prausnitzi*. *P<0.05; **P<0.01; ***P<0.001.

### Metagenomics functional prediction of the gut microbiome by response in the soluble HF group and *in vitro* functional effects of short chain fatty acids

Functional prediction at KEGG (Kyoto Encyclopedia of Genes and Genomes) pathway level 2 revealed that the gut microbiome in responders was enriched for carbohydrate metabolism pathways and in non-responders for membrane transport pathways (Figure 5F). For carbohydrate metabolism, the pathway with the highest level of enrichment, statistical analysis showed that responders had a significantly higher level than non-responders (Figure 5G). Short chain fatty acids (SCFAs), including acetate, propionate and butyrate, are major products of fibre fermentation by the gut microbiota. We showed that all three SCFAs increased histone acetylation (p=0.014 for 10 mM acetate, p=0.004 for 10 mM propionate, p<0.001 for 10 mM butyrate; Figure 5H) and tend to increase radiosensitivity (p=ns for acetate, p=ns for propionate, p=0.002 for butyrate in 8Gy; Figure 5I) of RT112 bladder cancer cells. Single SCFAs reduced cell proliferation (Figure 5J), while a physiological SCFA mixture conferred a stronger phenotype (Figure 5J, purple bar) which was shown in a time-dependent pattern as well (Additional file 1: Figure S5A).

To validate the anti-tumoural effects of *B. acidifaciens*, we treated the bladder tumour cells with bacterial supernatants of *B. acidifaciens* and its cross-feeding with *F. prausnitzii*, and compared their effects with *Bifidobacterium* (acetate-producer) and *F. prausnitzii* (butyrate-producer). Bacterial supernatants of *B. acidifaciens* and its cross-feeding with *F. prausnitzii* significantly increased cytotoxicity of bladder tumour cells compared to the other supernatants in day 2 (Figure 5K) and in day 3 (Additional file 1: Figure S5B).

### Correlation between the abundance of *B. acidifacien*s or *Parabacteroides* genus and mouse survival time in IR and non-IR cohorts

As *B. acidifaciens* was the ‘top hit’ for responders and the *Parabacteroides* genus was one of the top two ‘hits’ for non-responders in the soluble HF group, we explored how specific bacterial taxa affected mouse survival time. The correlation between *B. acidifaciens* abundance and time to culling was investigated across the diet groups. Some mice in the non-IR cohort lived as long as those in the IR cohort, ie. >40 days, which may be a reflection of 6 Gy being a relatively low dose of radiation. In the IR cohort, the time of culling positively correlated with *B. acidifaciens* abundance (R^2^=0.528, p<0.001). However, a similar correlation was not seen in the non-IR cohort (R^2^=0.085, p=0.357) (Additional file 1: Figure S6A). Using the time for tumours to treble in volume as the outcome measure, mice with high *B. acidifaciens* abundance had a significantly prolonged median survival time (Log-rank test: p<0.001) (Figure 6A). A similar finding was seen in the IR cohort (p=0.003), but not in the non-IR cohort (p=0.236). In the IR cohort, the time to culling negatively correlated with the abundance of *Parabacteroides* genus (Additional file 1: Figure S6B; R^2^=0.164, p=0.022). However, a similar correlation was not seen in the non-IR cohort (R^2^=0.084, p=0.360). Mice in the low *Parabacteroides* genus abundance group had no significant difference in median time to treble in volume compared to the high abundance group (Log-rank test: p=0.374) (Figure 6B). *B. acidifaciens* (p=0.200 for Kruskal-Wallis test) and *Parabacteroides* genus (p=0.005 for Kruskal-Wallis test) abundance was evenly distributed among all cages, which suggested that the existence of these taxa in the gut microbiome was not a cage-specific effect (Additional file 1: Figure S7 and Additional file 1: Table S2).

**Figure 6.**
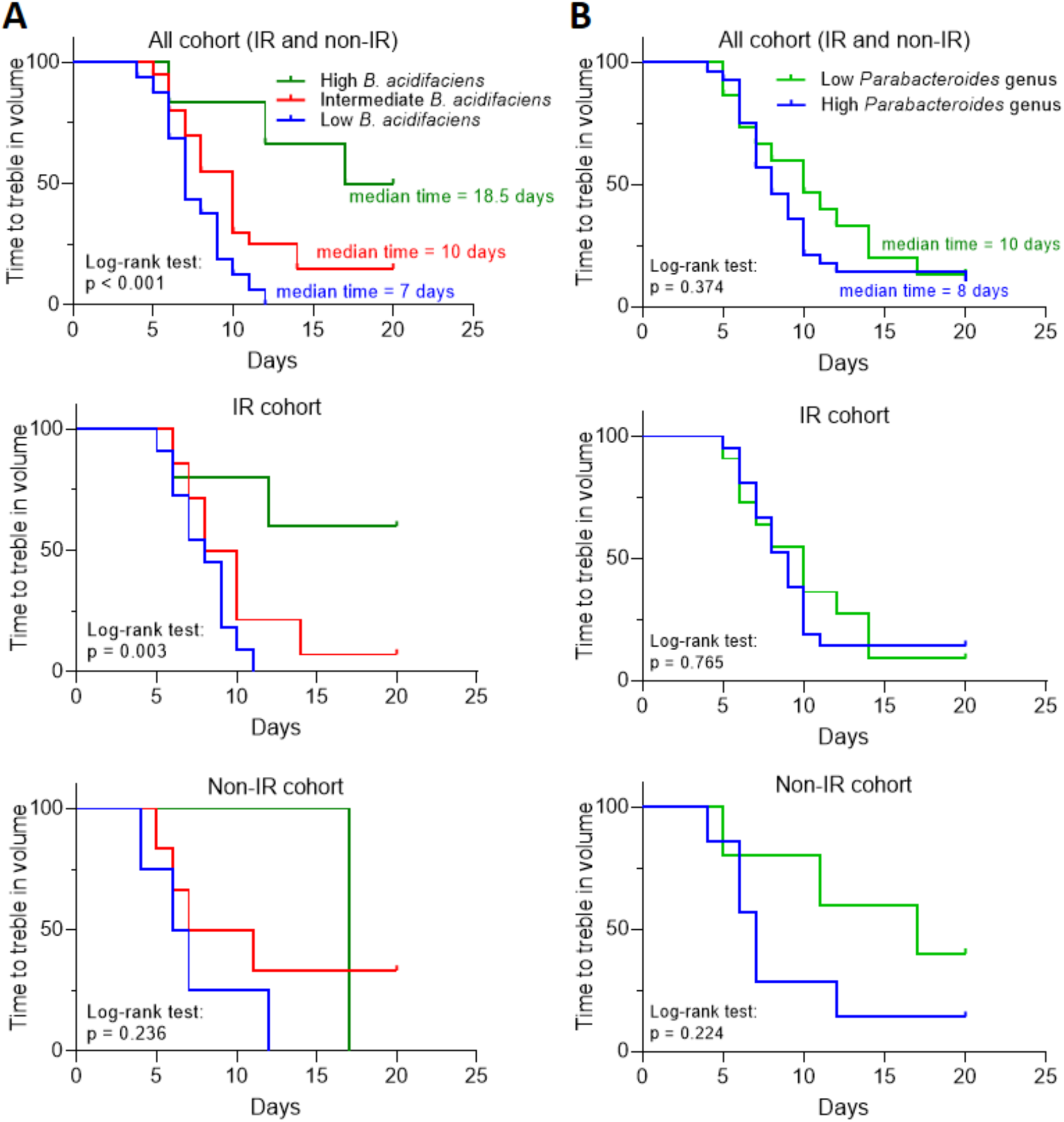
Abundance of OTUs within the gut microbiome is predictive of response to ionising irradiation. Kaplan-Meier (KM) plots of time for tumours to treble in volume, whole cohorts, IR cohorts, non-IR cohorts, based on (A) *B. acidifaciens* or (B) *Parabacteroides* genus abundance in IR and Non-IR cohorts from different diet groups combined. Comparison KM plots by log-rank test in mice with high abundance (green; relative abundance > 0.1), intermediate abundance (red; 0.1 > relative abundance > 0.01), or low abundance (blue; 0.01 > relative abundance) of *B. acidifaciens* in all, IR and non-IR cohorts. For *Parabacteroides* genus, relative abundance more than or equal to 0.01 was classified as high (blue), while less than 0.01 was classified as low (green).

## Discussion

To date, the literature studying the effects of dietary fibre intake or manipulation of the gut microbiota on tumour growth is limited. Wei *et al* showed that dietary fibre and the associated butyrate production reduced subcutaneous lymphoma tumour growth with associated upregulation of histone 3 acetylation (5). Hardman *et al* found slower growth of breast cancer xenografts in mice fed with fish oil concentrate (14). In contrast, Cougnoux *et al* found that Colibactin-producing *E. coli* enhanced tumour growth of colon cancer xenografts (15).

In this study, faecal and caecal microbiomes were investigated in mice fed the following: low fibre diet, low fibre diet with butyrate, high mixed fibre diet, high insoluble fibre (cellulose) diet and high soluble fibre (inulin) diet, and profiles of both microbiomes were correlated. The gut microbiomes were shaped by the different modified diets within two weeks and homogeneous gut microbiomes were seen in samples within groups. A distinct bacterial taxon was seen in each group: enrichment of *B. acidifaciens* in soluble HF, *Parabacteroides* in LF, *Akkermansia muciniphila* in LF with butyrate, and *Lachnospiraceae* in mixed HF.

The tumours were irradiated when they reached a volume of 50 mm^3^ (with no effect on diet on the time to reach this point) and monitored until they reached 350 mm^3^. During the course of the study, all mice developed an increased relative abundance of S24-7 family bacteria, thus indicating that their gut microbiomes had altered over the tumour growth period. Although the gut microbiomes became more heterogeneous, a notable cluster effect still existed in samples within groups. Mice responding to radiation in the soluble HF group were enriched with *B. acidifaciens* and non-responding mice were enriched with the *Parabacteroides* genus.

A predictive metagenomics study of the gut microbiome in responders was enriched for carbohydrate metabolism pathways. This implies a higher level of fibre fermentation occurring in responders, reflecting the selection of bacteria more able to ferment carbohydrates. This could result in more SCFA production in the faeces. Both butyrate and propionate have been proposed to increase histone deacetylase inhibition (16), which is a known mechanism of cellular radiosensitisation (17, 18). In this study, we demonstrated that these phenotypes exist in bladder cancer cells (Figure 5H and 5I). Furthermore, *in vitro* studies showed that SCFA reduced cell proliferation in liver cancer (19), induced apoptosis in lung cancer (20), and did both in breast cancer (21). Vecchia *et al*. found that acetate and propionate potentiated the anti-cancer effects of butyrate in leukemic, cervical adenocarcinoma, melanoma and breast cancer cells (22). We demonstrated similar findings in bladder cancer cells in this study (Figure 5J and Additional file 1: Figure S5A). Altogether, these data are supportive of carbohydrate metabolism or SCFA production being important in enhancing anti-cancer effects, including radiosensitisation.

When the mice from different diet groups were pooled, *B. acidifaciens* abundance was positively correlated with survival time, and mice with high *B. acidifaciens* had the longest median survival times using Kaplan-Meier survival analysis. Bacteroidetes, including *B. acidifaciens*, have been proposed to produce the metabolic end products acetate, succinate and possibly propionate, but not butyrate [15, 16]. High acetate levels could act as a substrate for butyrate production, given that acetate is necessary for butyrate production, particularly in the butyryl-CoA:acetate CoA-transferase pathway (23, 24). We speculate that the faecal butyrate levels could have been enhanced due to cross-feeding of butyrate-producing bacteria by *B. acidifaciens*. Proof-of-concept was previously demonstrated for this by cross-feeding butyrate-producing bacteria (*Faecalibacterium prausnitzii*) and acetate-producing bacteria (*Bifidobacterium adolescentis*) (25). Of note, Ramirez-Farias *et al* showed that inulin increased both *Faecalibacterium prausnitzii* and *Bifidobacterium adolescentis* in a human volunteer study (26).

A modified gut microbiota can augment the efficacy of anti-tumoural treatment. However, most studies to date are limited to chemotherapy and immunotherapy, reviewed in (27, 28). Recently, Herranz *et al* found that depletion of gram positive bacteria in the gut by vancomycin enhanced radiotherapy-induced anti-tumour immune response and further delayed tumour growth. However, the authors gave very large 21 Gy single fraction irradiation doses. Furthermore, reductions in abundance/absence of gram negative bacteria (including *Bacteroides* and *S24-7*) were also seen, with increased *Parabacteroides*, which could be of significance. Here we only gave a 6 Gy single fraction of IR which is more clinically relevant, and our findings were not immunologically-mediated via T cells, as CD1 nude mice lack T cells. To our knowledge, ours is the first study to provide evidence that a high soluble fibre diet, and its subsequent modification of the gut microbiome, can act to radiosensitise tumours.

Furthermore, *B. acidifaciens* was identified as a potential radiosensitiser because its abundance was enhanced by a high soluble fibre diet and positively correlated with tumour response to radiation and survival time in the IR cohort. This bacterium was first isolated in 2000 and was so named because it reduces the pH level of peptone-yeast broth with Fildes’ digest (13). Consistent with our findings, Marques *et al* demonstrated that a high-fibre diet markedly increased the prevalence of *B. acidifaciens* (29). Another study showed *B. acidifaciens* to be enriched in normal human subjects, compared to patients with inflammatory bowel disease (30), and *B. acidifaciens* increased insulin sensitivity and prevented further obesity (31). However, the effect of this bacteria on tumour growth is still controversial. A study found increased *B. acidifaciens* abundance associated with hepatocellular carcinoma induced by a streptozotocin-high fat diet (32). In contrast, *B. acidifaciens* reduces the isoflavone genisten, which is associated with increased risk of breast cancer (33) and *B. acidifaciens* was shown to contribute to the anti-tumour effect of medicinal Gynostemma saponins (34). In this study, we revealed that bacterial supernatant from *B. acidifaciens* and its cross-feeding with *F. prausnitzii* caused significantly higher levels of cytotoxicity compared to the other supernatants (Figure 5K and Additional file 1: Figure S5B). This result supports our finding that *B. acidifaciens* may drive the radiosensitising effect. Moreover, *B. acidifaciens in vitro* has a greater effect on cell kill than *F. prausnitzii* (butyrate-producer; p<0.001), implying that metabolites other than butyrate may be involved in its effect. Studies suggest that a broad range of gut microbiota-derived metabolites can enhance anti-tumoural effects or tumour response to anti-cancer treatments (35-37), so future work should include undertaking a global metabolomic analysis of *B. acidifaciens*-produced metabolites to identify radiosensitisers other than SCFAs with similar profiles.

We also found *Bacteroidales S24-7* (or *Candidatus Homeothermaceae* (38) or *Muribaculaceae* (39)), an uncultured bacterium with limited characterisation, to be highly abundant in our study, especially in the mixed HF and soluble HF group. The prevalence of this bacterium in humans is 20% (40). However, increased prevalence of *Bacteroidales S24-7* of up to 70% has been shown in the Hadza tribe of Tanzania who consume tubers containing large amounts of soluble fibre (40). This indicates that high abundance of *Bacteroidales S24-7* found in this study when the tumours reached 350 mm^3^ might be induced by soluble high fibre.

Environmental contamination is an inevitable issue in microbiome studies (41). To minimise the influence of contamination in this study, bacterial loads of samples were quantified and appropriate negative controls were included. Bacterial loads from luminal contents and tissue samples contained more than 10^4^ CFUs which overrode the environmental bacteria communities. Furthermore, the bacterial compositions of the PBS negative controls were very different from those of the study groups, so the environmental microbiome was considered not to be a major source of bias in this study.

Although a strong correlation between *B. acidifaciens* abundance and tumour response to irradiation was seen, a limitation of this study is that only one cohort of mice was studied. Further studies are needed to determine the causal relationship between *B. acidifaciens* and radiosensitisation, and its underlying aetiology. This could be achieved by oral gavage of *B. acidifaciens* with or without other bacteria in gnotobiotic mice and *in vitro* culture studies of *B. acidifaciens*. As both gut bacteria (42) and irradiation (43) directly interact with the immune system, further *in vivo* studies need to be conducted in an immune competent model to reveal how immunomodulation might contribute to the radiosensitisation.

## Conclusion

A high soluble fibre diet increased responses of RT112 subcutaneous xenografts in CD1 nude mice to ionising radiation and this phenotype was associated with higher relative abundance of *B. acidifaciens*. Possible mechanisms mediating this effect, which require further investigation, include: (1) increased concentrations of metabolites, including butyrate or other short-chain fatty acids in tumours, acting via HDAC inhibition or via other pathways; (2) suppression of overgrowth of unfavourable bacteria, such as *Parabacteroides* genus; and/or (3) enhancement of anti-tumoural immunity. Our findings suggest that dietary fibre modification and the resultant modification of the gut microbiome might be exploited to improve tumour responses to radiotherapy in human patients.

## Methods

### Mice and mouse diets

All animal work was done in accordance with UK Home Office Guidelines, following the ARRIVE (Animal Research: Reporting of In Vivo Experiments) guidelines, and approved by the University of Oxford Animal Welfare and Ethical Review Body (AWERB), under University of Oxford project licences P4B738A3B and P8484EDAE. Group sizes were chosen to detect large effect sizes by using a G-Power analysis program. All mice were purchased from Charles Rivers UK Ltd.

CD1-nude female mice at 6-7 weeks old were housed in a temperature-controlled environment with a 12-h reversed-phase light/dark cycle (lights on 21:00 h) and provided with food and water *ad libitum*. These mice are immunodeficient, lacking a thymus and therefore unable to produce T-cells. Mice were randomised in Excel using the RAND function into four groups. At 7 to 8 weeks of age, mice were injected subcutaneously with RT112 bladder cancer cells (DSMZ, Germany) and started receiving either a low fibre diet (2 g cellulose/3850 kcal), a high insoluble fibre diet (100 g cellulose/3850 kcal), a high soluble fibre diet (100 g inulin/3850 kcal) or a high mixed fibre diet (50 g cellulose + 50 g inulin / 3850 kcal) for a maximum time of 9 weeks or until they were culled when the tumours reached 350 mm^3^. Faeces, caecal contents, and proximal and distal colons from the first cohort were taken when the tumour reached 50 mm^3^ (each group n=3) without irradiation to investigate the microbiome at baseline. Faeces and caecal contents from the second cohort were taken when the tumour reached 350 mm^3^ after IR (each group n=8) or without IR (each group n=3) or at the end of study (9 weeks after xenograft) to study the association between the gut microbiome composition and tumour response.

### Xenograft model and irradiation method

Mice were injected subcutaneously under anaesthesia into the right flank with 5 × 10^6^ human bladder cancer cells (RT112) in RPMI medium (Sigma Aldrich) with phenol red-free Matrigel (BD Biosciences) at a total volume of 100 μl (1:1 ratio cell suspension to Matrigel). Tumour growth was measured three times a week and size determined by calipers using (length x width x height x Π/6). To assess the effects of different dietary fibres on tumour growth after irradiation *in vivo*, mice received ionising radiation to the tumour (6 Gy, single fraction, 300 kV, using a Gulmay-320 cabinet irradiator, Xstrahl Inc, UK). A dose of 6 Gy was chosen due to the promising effect in our previous radiosensitisation experiments with a HDAC inhibitor (manuscript in preparation.)

### Microbiome sample collection and DNA extraction

All samples were transported on ice and kept at −20°C for less than 2 hours before DNA extraction. Bacterial genomic DNA was extracted using a DNeasy PowerSoil DNA Isolation Kit (QIAGEN Ltd, Manchester, UK), as per the Human Microbiome Project (44). Briefly, by adding sodium dodecyl sulfate (SDS), microbial cells were lysed by mechanical disruption with a ceramic bead set on 3,000 rpm for 10 minutes, followed by binding of DNA tightly to a silica membrane in a Spin Filter device at high salt concentrations. Eventually, DNA was collected into sterile elution buffer and quantified using a NanoDrop spectrophotometer. All DNA samples were kept at −80°C. All samples were collected and handled in sterile containers and equipment to minimise contamination. Those sent for sequencing (Omega Bioservices, Georgia, USA) were dried in an Eppendorf concentrator 5301 (Eppendorf North America Inc, USA) at a rotational speed of 1,400 rpm and centrifugal force of 240 x g for 1 hour at 30°C.

### Faecal butyrate levels quantification

Faecal samples were first homogenised in ice cold Millipore Synergy purified water. Thereafter, 20 μL of sample or standard was taken and 10 μL of internal standard (valeric acid, Alfa Aesar, UK) added prior to the addition of 5 μL 15% percholoric acid. Samples were mixed and centrifuged at 12,000 g for 15 min at 4°C followed by direct injection (10 μL) of the supernatant. High-performance liquid chromatography (HPLC) separation was carried out using a Waters Acquity H-Class Quarternary Solvent Manager with mobile phases of 0.1% formic acid in water (A) and methanol (B) and a gradient of 35-75% B on a Waters Acquity CSH C18, 1.7 μm, 100 × 2.1 column. Butyrate and internal standard (IS) were detected by mass spectrometry with a Waters Acquity TQ detector in positive electrospray ionisation mode. Butyrate was detected with a cone voltage of 20 V at selected ion recording (SIR) of *m/z* 88.41 (M+H) and IS with a cone voltage of 15 V and SIR of *m/z* 103.2 (M+H).

### Cell line, drugs, and irradiator

The RT112 bladder carcinoma cell line was obtained from the American Type Culture Collection. This cell line was cultured inRPMI-1640 medium (Sigma), supplemented with 10% fetal bovine serum (Invitrogen). Mycoplasma testing was negative. Sodium acetate, sodium propionate and sodium butyrate were purchased from Sigma-Aldrich (Gillingham, United Kingdom) and used in dH_2_O. Cells were irradiated in complete medium at a dose rate of 1.5 Gy/min using a Gamma-Service Medical GmbH GSR D1 irradiator.

### Bacterial strain and its supernatant

All bacterial strains were obtained from DSMZ-German collection of microorganisms. Three strains of bacteria, namely *B. acidifaciens* (*BA*; DSM 15896), *Bifidobacterium animali*s (*Bif*; DSM10140), *F. prausnitzii* (*FP*; DSM17677), and two cross-feeding combinations (*BA+FP* and *Bif+FP*) were cultured in Gifu Anaerobic Broth, Modified (GAM; Nissui Pharmaceutical, Japan). Medium broth was prepared anaerobically in a Coy anaerobic chamber (Coylabs, US, N2 95%, H2 5%) and sealed into 10 mL glass tubes before being autoclaved at 121°C for 15 minutes. Bacteria (10^6^ CFU/mL starting population for each strain) were inoculated into these sealed tubes ascetically by injecting through a needle. They were then placed in a 37°C incubator for 24 hours with constant shaking. Samples were centrifuged at 4000 × g for 10 minutes to remove all particulate matter. Supernatants were then filtered through a 0.22 μm polyethersulfone syringe filter (Millipore).

### Colony formation assay

Cells were seeded in 6-well plates at appropriate densities in triplicate, treated with short-chain fatty acids, namely acetate, propionate and butyrate, at appropriate concentrations for 24 hours, and irradiated with 0, 2, 4, 6, and 8 Gy. After culturing for 10 days, colonies were fixed and stained with 0.5% crystal violet in dH_2_O and 20% methanol for 5 minutes. Finally, they were quantified using a GelCount colony counter (Oxford Optronix). The surviving fraction was calculated by normalising the number of colonies for each condition to the unirradiated control.

### Cell survival analysis

MTT 3-(4,5-dimethylthiazol-2-yl)-2,5-diphenyltetrazolium bromide assay was applied to assess cell viability by adding 0.25 mg/mL MTT (Life Technologies) to cells at 37 °C for 1 h in the end of experiment. The absorbance at 595 nm of MTT-formazan was detected spectrophotometrically using an POLARstar Omega Microplate Readers (BMG Labtech) after dissolution of the crystals in isopropanol. The percentage of cell viability was calculated by the formula: [Experimental group / Control group] x 100%.

### Western blots

Western blot samples were prepared as described in (45). Protein was visualised using the following antibodies: H3K18Ac (Cell Signaling Technology, #9675), H3 (Cell Signaling Technologies, #4499S), and β-actin (Abcam, #A1978), and an infrared LiCor Odyssey imaging system (LiCor Biosciences).

### Identification and quantification of bacterial DNA

The microbiota of the contents of the intestinal tracts and the intestinal wall of the proximal and distal colon (tissue) was quantified by PCR of 16S rRNA. This was performed on genomic DNA extracted as described above. The PCR was performed using primers - V3F (CCAGACTCCTACGGGAGGCAG) and V3R (CGTATTACCGCGGCTGCTG) (46). All primers were purchased from Sigma. For each sample, Phire Tissue Direct PCR Master Mix (Thermo Fisher Scientific) was used to amplify the 16S rRNA gene hypervariable V3 region (product size = 200 bp). PCR amplifications were performed using the following conditions: 98°C for 5 minutes followed by 35 cycles at 98°C for 5 seconds each, 66.3°C for 5 seconds, and 72°C for 30 seconds and a final extension step at 72°C for 1 minute. The amplification products were visualised on a 1% agarose gel after electrophoretic migration of 5 μl of amplified material. A standard curve was created from serial dilutions of *Escherichia Coli* from 1 × 10^2^, 1 × 10^4^, 1 x10^6^ colony-forming units (CFU) which was quantified by CFU assay. All samples were run in duplicate. In CFU assay, 20 μL of serial dilution of *E. Coli* was incubated onto Luria-Bertani (LB) agar plates, and colonies were counted and bacterial concentrations of the original samples were estimated after 24 hours incubation.

### Bacterial 16S rRNA gene sequencing

16S rRNA gene sequencing methods were adapted from the methods developed for the NIH-Human Microbiome Project (44, 47). Raw 16S rRNA reads and metadata have been made available in Figshare (https://figshare.com/projects/The_gut_microbiota_may_drive_the_radiosensitising_effect_of_a_high_fibre_diet/68393) (48). The amplification and sequencing of 16S rRNA gene V3V4 region were done commercially by Omega Bioservices (Georgia, USA) on a MiSeq platform (Illumina, Inc, San Diego, CA) using the 2×300 bp paired-end protocol, yielding paired-end reads with near-complete overlap. The primers containing adapters for Miseq sequencing were used for amplification and single-end barcodes, allowing pooling and direct sequencing of PCR products (49). PBS negative controls were included to eliminate the confounding effects of environmental contamination. All 16S rRNA gene-based metagenomic analysis was conducted using a QIIME2 platform (50). Quality filtered sequences with >97% identity were clustered into bins known as Operational Taxonomic Units (OTUs), using open-reference OTU picking. The relative abundance of each OTU was obtained from all samples. In the taxonomic analysis, the microbiome at the phylum, class, order, family, genus and species levels was classified with reference to the Greengenes database (51).

The analysis pipeline was as follows:

i. All sequences were trimmed to a length of 240, since the quality dropped above this length based on the sequence quality plots.
ii. De-noised sequencing errors by using the “Deblur” plugin in QIIME2 (52).
iii. Taxonomic assignment was performed with Greengenes (53) by the “feature-classifier” command.
iv. To visualise the differences in microbial composition between gut contents and tissue, a taxonomic profile was generated by conducting differential abundance analysis using balances in gneiss.
v. To identify the features characterising the differences between groups, the LEfSe method of analysis was performed to compare abundances of all bacterial clades (54). By validation using the Kruskal-Wallis test at the α setting of 0.05, effect size was obtained by LDA (linear discriminant analysis) based on the significantly different vectors resulting from the comparison of abundances between groups.
vi. To validate the significance of enrichment of bacterial taxa among different groups, discrete false-discovery rates (DS-FDR) were calculated (55)
vii. A phylogenetic tree was generated by using the “phylogeny” plugin in QIIME2.
viii. To investigate the alpha and beta diversity, the diversity commands of “alpha-group-significance” and “beta-group-significance” were used to obtain Shannon’s index, and Bray-Curtis dissimilarity. A principal coordinates (PCoA) plot was obtained by using the Emperor Tool based on the results of Bray-Curtis dissimilarities.
ix. The OTU table was rarefied using the “alpha-rarefraction” command in QIIME2. The alpha rarefraction plot showed the richness of the samples with increasing sequence count.
x. To predict the metagenome functional profiles, PICRUSt, a bioinformatics software package, was used to collapse predicted functions (KEGG Orthology; KO) based on 16S rRNA surveys into higher categories (KEGG pathway) after picking OTUs and normalisation (56).

### Statistics

Power calculations for the number of mice per group were done using G*Power software version 3.1.9.4 (57). Alpha diversity and relative abundance of specific bacterial taxa were compared using the Kruskal-Wallis test following by Dunn’s multiple comparison test. All mice were classified into high, intermediate or low diversity groups based on tertiles of distribution. Time to treble in volume was defined as the interval (in days) from the date of irradiation (growth to 50 mm^3^) to the date for the tumour to treble in volume. Tumour growth curves were analysed for each group, and their slopes were compared using one-way ANOVA. The LEfSe method of analysis was applied to determine the difference in bacterial taxa, using the Kruskal-Wallis test. Significantly different taxa presented from the previous comparison applying LEfSe method were used as input for LDA, which produced an LDA score. Volcano plots showed the significance of the taxa which are different among different groups, with log_10_ (FDR-adjusted p-values) on the y-axis and median-adjusted effect sizes on the x-axis. In addition, mice were also classified as having high, intermediate and low abundance of *B. acidifaciens* or high and low abundance of *parabacteroides* genus based on the relative abundance of these taxa in the gut microbiome sample. All analyses were conducted in QIIME2 and GraphPad Prism version 8.0 (La Jolla, CA). All data in *in vitro* studies are representative of 3 independent biological replicates unless otherwise stated, with results shown as mean and standard deviations. One-way ANOVA with Dunnett’s multiple comparison test was performed to analyse the data of Western Blots and MTT assays. Two-way ANOVA with Dunnett’s multiple comparison test was used to analyse the linear quadratic survival curves in colony formation assay.

## List of abbreviations

ABC: ATP-binding cassette
*B. acidifaciens*: Bacteroides acidifaciens
BBN: N-butyl-N-(4-hydroxybutyl)-nitrosamine
CFU: colony formation unit
E. Coli: Escherichia coli
FDA: false-discovery rate
Gy: Gray
HAT: histone acetyltransferase
HDAC: histone deacetylase
HF: high fibre
Insoluble HF: insoluble high fibre
IR: irradiation
KEGG: Kyoto Encyclopedia of Genes and Genomes
LC-MS: liquid chromatography–mass spectrometry
LDA: linear discriminant analysis
LEFSe: linear discriminant analysis effect size
LF: low fibre
LF+B: low fibre plus butyrate
MIBC: muscle-invasive bladder cancer
Mixed HF: mixed high fibre
MTT: 3-(4,5-dimethylthiazol-2-yl)-2,5-diphenyltetrazolium bromide
OTU: operational taxonomic unit
PCoA: principal coordinates analysis
PICRUSt: Phylogenetic Investigation of Communities by Reconstruction of Unobserved States
SCFA: short chain fatty acid
SDS: sodium dodecyl sulfate
Soluble HF: soluble high fibre

## Acknowledgements

We thank Professor Simon Kroll and Dr Anderson Ryan for their very helpful comments. We thank Dr Jia-Yu Ke at Research Diets, Inc. for formulation of the mouse diets, Dr Lisa Folkes for assistance with the faecal butyrate quantification and Omega Bioservices (Georgia, USA) for the 16S rRNA gene sequencing on a MiSeq platform.

## Declarations

### Funding

This work was funded by Cancer Research UK Programme grant C5255/A15935 and Wellcome Trust Investigator Award 209397/Z/17/Z. The funding body had no role in the design of the study, collection, analysis, interpretation of data or in writing the manuscript. **Availability of data and materials**

The datasets generated and/or analysed during the current study are available in the Figshare repository, https://figshare.com/projects/The_gut_microbiota_may_drive_the_radiosensitising_effect_of_a_high_fibre_diet/68393 (48).

### Authors’ contributions

CKT extracted the DNA from mouse samples, performed the analysis and interpretation of the data and drafted the manuscript. SP performed the animal experiments, collected the faeces, caecal contents, intestinal tissue and blood. XW produced the bacterial supernatants. AH measured the faecal butyrate levels. AEK conceived the study, supervised the work, and revised the manuscript. All authors read and approved the final manuscript.

### Ethics Approval and Consent to Participate

All animal protocols were approved by the University of Oxford Clinical Medicine Animal Welfare Ethics Review Board and conducted under animal project licences (PPL) P4B738A3B and P8484EDAE.

## Additional files legends

Additional file 1: FigS1 - Similar bacterial components in the faecal and caecal microbiomes. FigS2 - Faecal butyrate levels and time taken for tumours to reach 50 mm^3^. FigS3 - Differences in composition of the gut microbiome when tumours reached 350 mm^3^. FigS4 - Individual mouse tumour growth curves. FigS5 - Cell survival analysis of RT112 bladder tumour cells treated with SCFAs and bacterial supernatants. FigS6 - Correlation of time to culling with *B. acidifaciens* or *Parabacteroides* genus abundance different groups. FigS7 - Effect of cage location of mice on relative abundance of *B. acidifaciens* and *Parabacteroides* genus.

