## Additional file 1 for "Association of *Bacteroides acidifaciens* relative abundance with high-fibre diet-associated radiosensitisation"

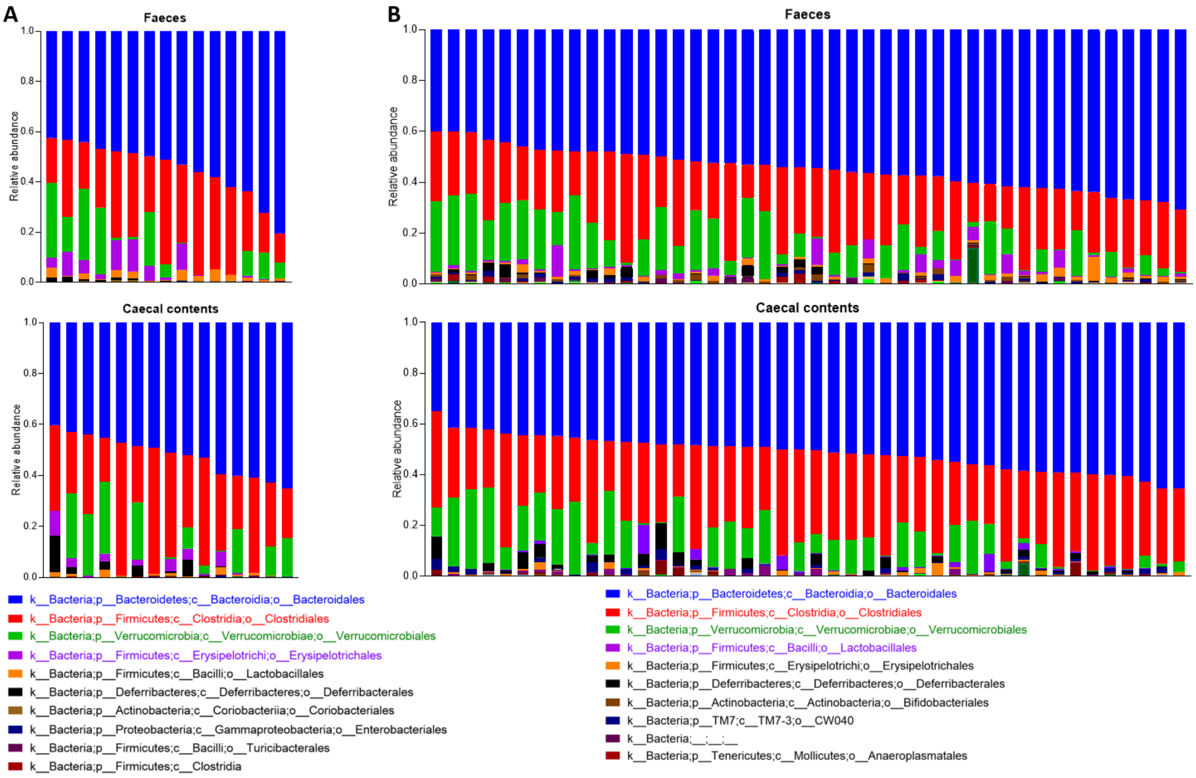


**Figure S1. Similar bacterial components in the faecal and caecal microbiomes.** Phylogenetic composition of faecal and caecal microbiomes at the order level when tumours reached (A) 50 $mm^3$ (n = 15) and (B) 350 $mm^3$ (n = 44). Samples were collected from faeces and caecal contents, and sorted by *Bacteroidales* proportion.

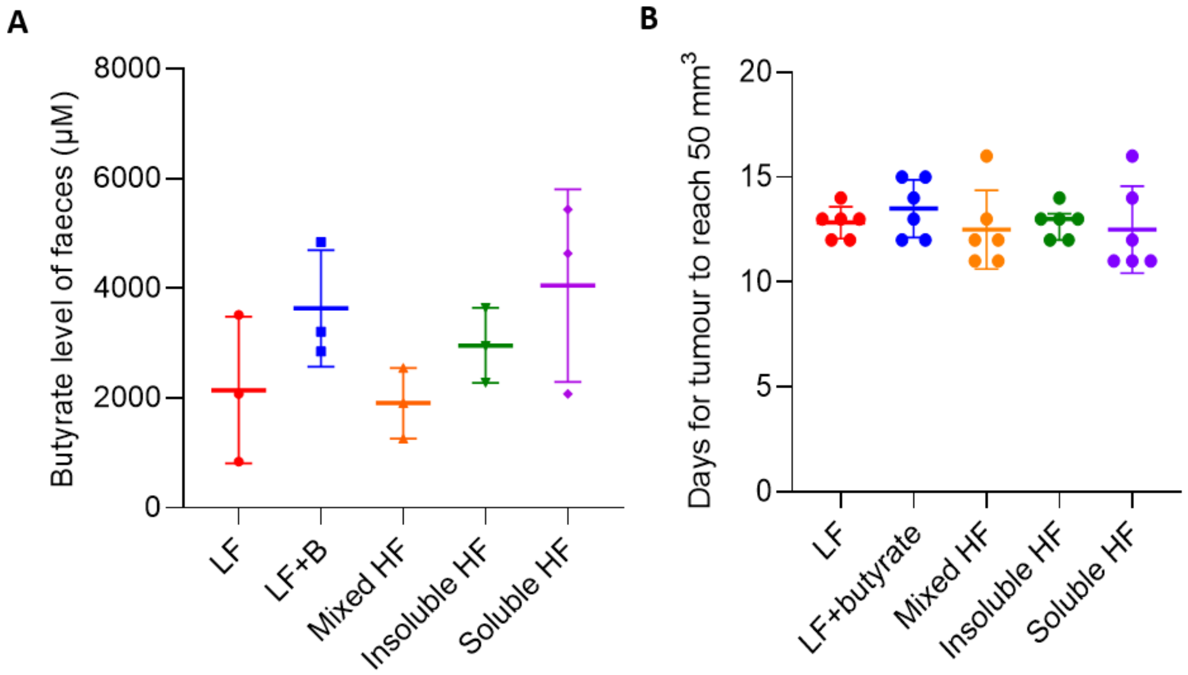


**Figure S2. Faecal butyrate levels and time taken for tumours to reach 50 mm$^3$.** (A) Butyrate levels in the faeces at the time of culling. (B) All mice were culled when tumours reached 50 mm$^3$, between 11 to 16 days after tumour inoculation, mean 12.8 (SD $\pm$ 1.4) days.

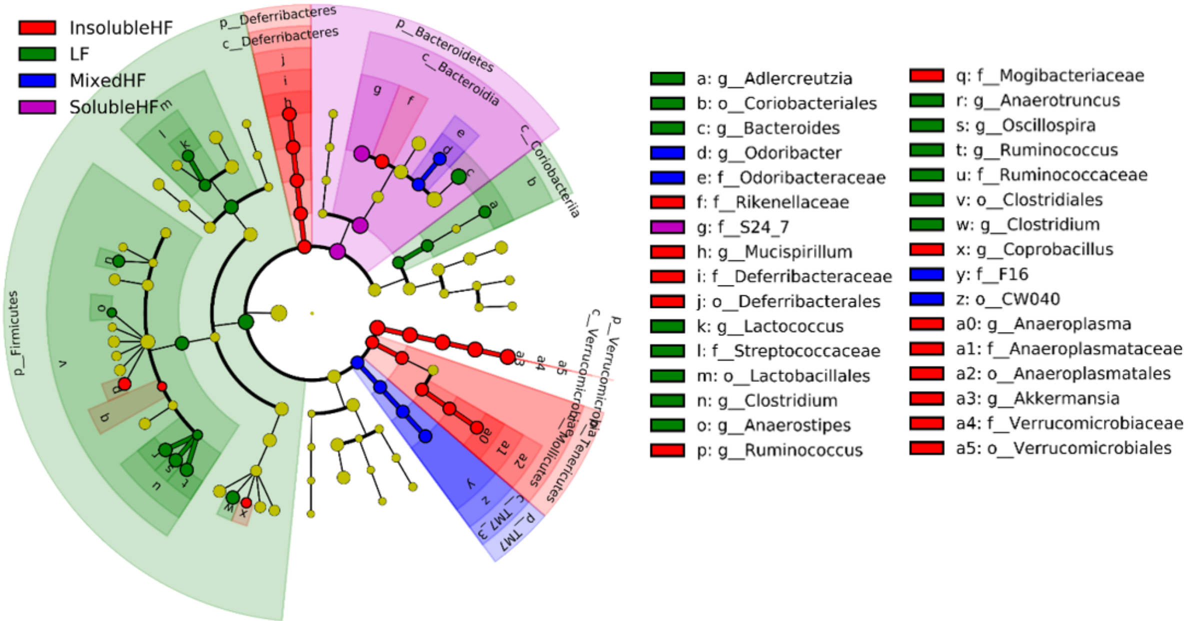


**Figure S3. Differences in composition of the gut microbiome when tumours reached 350 mm$^3$.** Taxonomic cladogram from LEfSe showing differences in bacterial taxa at the genus level in the IR cohort when the tumours reached 350 mm$^3$. The low fibre diet increased *Adlercreutzia, Coriobacteriales, Bacteroides, Lactococcus, Streptococcaceae, Lactobacillales, Oscillospira, Ruminococcus, Ruminococcaceae, Clostridiales, Clostridium*, the high mixed fibre increasing *Odoribacter, Odoribacteraceae, F16, CW040*, the high insoluble fibre diet increased *Rikenellaceae, Mucispirillum, Deferribacteraceae, Deferribacterales, Ruminococcus, Mogibacteriaceae, Coprobacillus, Anaeroplasma, Anaeroplasmataceae, Anaeroplasmatales, Akkermansia, Verrucomicrobiaceae, Verrucomicrobiales*, and the high soluble fibre diet increased *S24-7.*

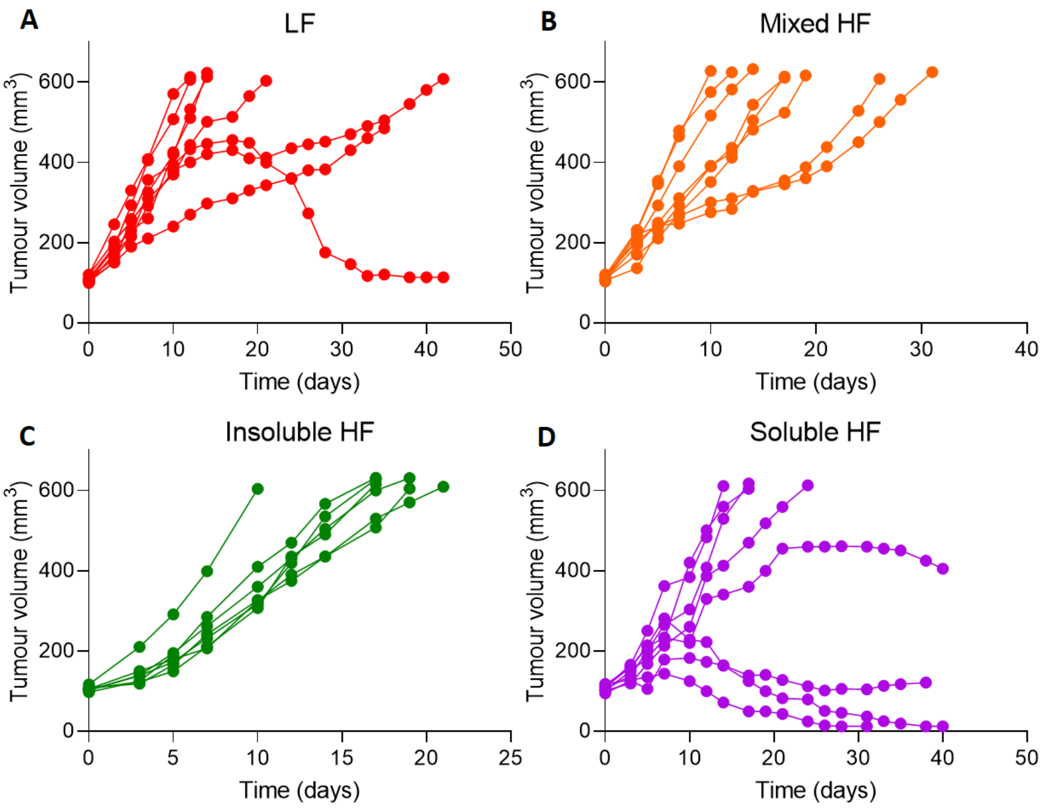


**Figure S4. Individual mouse tumour growth curves.** Tumour growth in RT112 flank xenografts irradiated with 6 Gy IR, in mice fed low fibre (A), high mixed fibre (B), high insoluble fibre (C) and high soluble fibre diets (D) (n = 8 for each group).

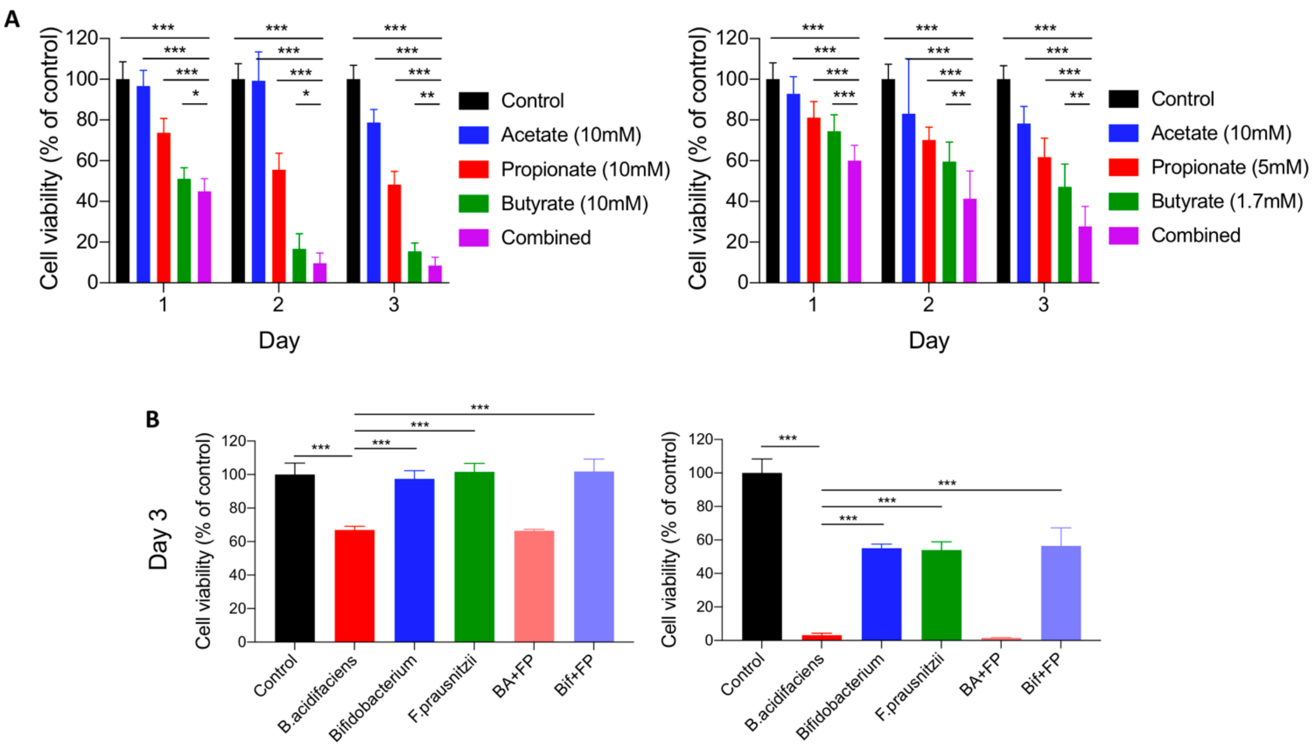


**Figure S5. Cell survival analysis of RT112 bladder tumour cells treated with SCFAs and bacterial supernatants.** (A) Inhibition of cell viability of RT112 cells single SCFA and combined SCFAs mixture in a time-dependent manner (N=3). The combined SCFAs denote the mixtures of 10 mM butyrate, 10 mM propionate, 10 mM butyrate for the left-hand graph and the mixtures of 10 mM butyrate, 5 mM propionate, 1.7 mM butyrate for the right-hand graph. (B) Reduced cell survival of RT112 cells by bacterial supernatants at day 3 (N=1). *BA+FP* denotes the cross-feeding of *B. acidifaciens* and *F. prausnitzii*, while *Bif+FP* denotes the cross-feeding of *Bifidobacterium* and *F. prausnitzi.* *P<0.05; **P<0.01; ***P<0.001.

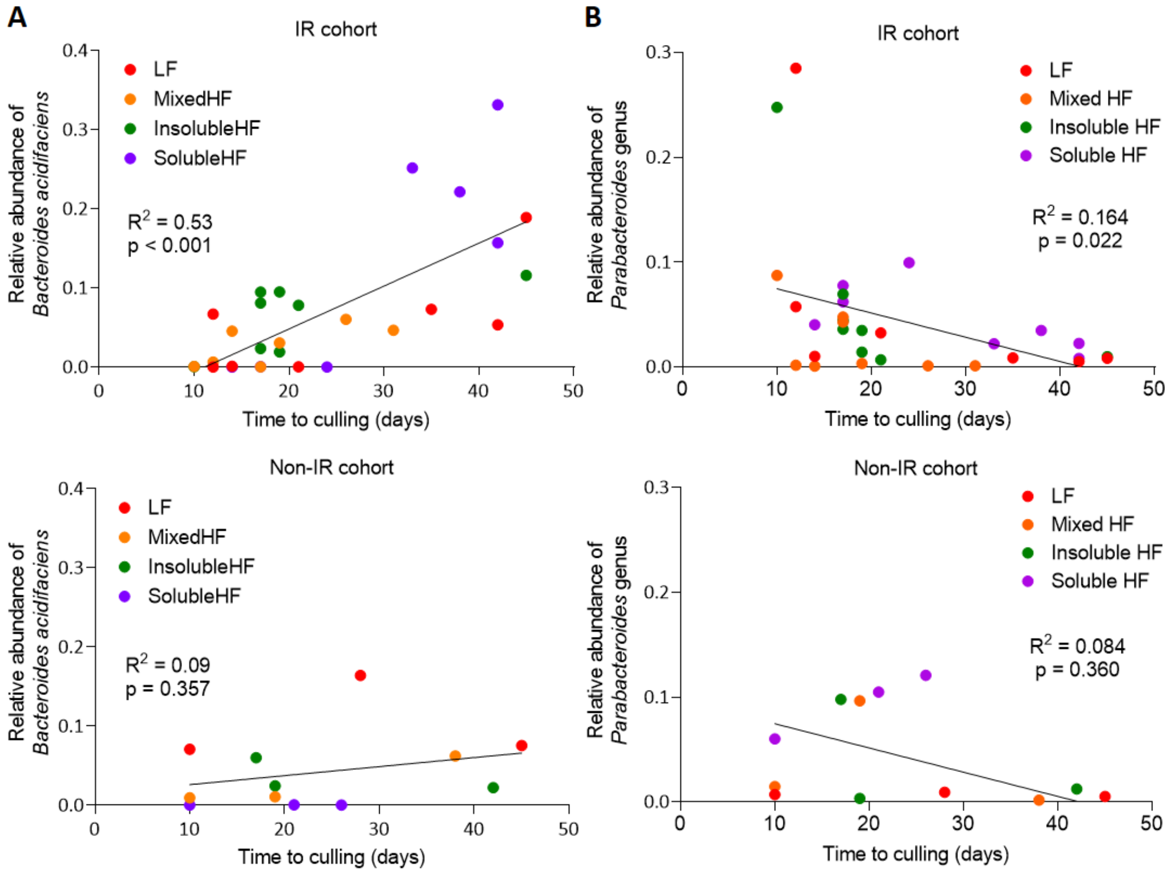


**Figure S6. Correlation of time to culling with *B. acidifaciens* or *Parabacteroides* genus abundance different groups.** (A) Correlation between time of culling versus (A) *B. acidifaciens* abundance (IR cohort, $R^2 = 0.53$, $p < 0.001$; Non-IR cohort, $R^2 = 0.09$, $P = 0.357$) or *Parabacteroides* genus abundance (IR cohort, $R^2 = 0.164$, $P = 0.022$; Non-IR cohort, $R^2 = 0.084$, $P = 0.360$) in the gut microbiome.

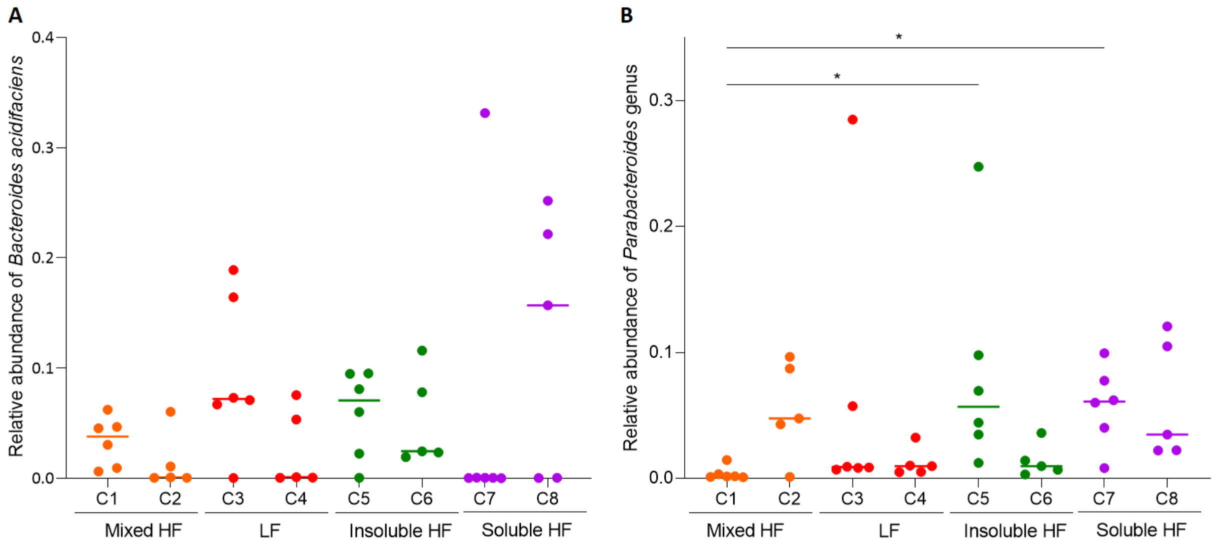


**Figure S7. Effect of cage location of mice on relative abundance of *B. acidifaciens* and *Parabacteroides* genus.** Comparison of (A) *B. acidifaciens* (p=0.200) or (B) *Parabacteroides* genus (p=0.005) abundance among mice from cage 1 (C1) to cage 8 (C8) by Kruskal-Wallis test.

**Table S1. Rodent diets used in the study with varying levels of cellulose or inulin per 4000 kcal.**

| | Low fibre | | High insoluble fibre | | High soluble fibre | | High mixed fibre | |
|---|---|---|---|---|---|---|---|---|
| | 2 gm Cellulose/3850kcal | | 100 gm Cellulose/3850kcal | | 100 gm Inulin/3850kcal | | 50 gm Cellulose + 50 gm Inulin/3850kcal | |
| | gm% | kcal% | gm% | kcal% | gm% | kcal% | gm% | kcal% |
| Protein | 21 | 20 | 19 | 20 | 20 | 20 | 20 | 20 |
| Carbohydrate | 67 | 64 | 61 | 64 | 59 | 60 | 60 | 62 |
| Fat | 7 | 16 | 7 | 16 | 7 | 16 | 7 | 16 |
| Total | | 100 | | 100 | | 96 | | 96 |
| kcal/gm | 4.20 | | 3.81 | | 3.95 | | 3.88 | |
| **Ingredient** | **gm** | **kcal** | **gm** | **kcal** | **gm** | **kcal** | **gm** | **kcal** |
| Casein | 200 | 800 | 200 | 800 | 200 | 800 | 200 | 800 |
| L-Cystein | 3 | 12 | 3 | 12 | 3 | 12 | 3 | 12 |
| Corn starch | 397.486 | 1590 | 397.486 | 1590 | 359.986 | 1440 | 378.736 | 1515 |
| Maltodextrin 10 | 132 | 528 | 132 | 528 | 132 | 528 | 132 | 528 |
| Sucrose | 100 | 400 | 100 | 400 | 100 | 400 | 100 | 400 |
| Cellulose, BM200 | 2 | 0 | 100 | 0 | 0 | 0 | 50 | 0 |
| Inulin | 0 | 0 | 0 | 0 | 100 | 0 | 50 | 75 |
| Soybean oil | 70 | 630 | 70 | 630 | 70 | 630 | 70 | 630 |
| t-Butylhydroquinone | 0.014 | 0 | 0.014 | 0 | 0.014 | 0 | 0.014 | 0 |
| Mineral mix S10022G | 35 | 0 | 35 | 0 | 35 | 0 | 35 | 0 |
| Vitamin mix V10037 | 10 | 40 | 10 | 40 | 10 | 40 | 10 | 40 |
| Choline bitartrate | 2.5 | 0 | 2.5 | 0 | 2.5 | 0 | 2.5 | 0 |
| **Total** | **952** | **4000** | **1050** | **4000** | **1012.5** | **4000** | **1031.25** | **4000** |
| Total Cellulose (gm/kg diet) | 2.1 | | 95.2 | | 0 | | 48.5 | |
| Inulin (gm/kg diet) | 0 | | 0 | | 98.8 | | 48.5 | |

### Table S2. Details mouse diets, cages, *B. acidifaciens* relative abundance and time of culling

| Diet | **Low fibre** | | | | | | | | | | |
|---|---|---|---|---|---|---|---|---|---|---|---|
| Cage | 3 | | | | | | 4 | | | | |
| ID | C3.3 | C3.1 | C3.10 | C3.30 | C3.NM | C3.4 | C4.10 | C4.1 | C4.NM | C4.30 | C4.3 |
| IR/Non-IR | Non-IR | IR | IR | IR | IR | Non-IR | Non-IR | IR | IR | IR | IR |
| *B.acidifaciens* abundance | 0.164 | 0.000 | 0.067 | 0.189 | 0.073 | 0.071 | 0.075 | 0.000 | 0.053 | 0.001 | 0.001 |
| *Parabacteroides* genus abundance | 0.009 | 0.285 | 0.057 | 0.008 | 0.009 | 0.007 | 0.005 | 0.032 | 0.005 | 0.010 | 0.010 |
| Time for tumour to treble in volume | 17 | 6 | 6 | 6 | 14 | 5 | - | 9 | 8 | 7 | 8 |
| Time to culling (days) | 28 | 12 | 12 | 45 | 35 | 10 | 45 | 21 | 42 | 14 | 14 |

| Diet | **Mixed high fibre** | | | | | | | | | | |
|---|---|---|---|---|---|---|---|---|---|---|---|
| Cage | 1 | | | | | | 2 | | | | |
| ID | C1.NM | C1.10 | C1.3 | C1.1 | C1.30 | C1.4 | C2.10 | C2.NM | C2.3 | C2.30 | C2.1 |
| IR/Non-IR | IR | IR | IR | Non-IR | Non-IR | IR | IR | IR | IR | Non-IR | IR |
| *B.acidifaciens* abundance | 0.045 | 0.030 | 0.047 | 0.009 | 0.062 | 0.006 | 0.001 | 0.000 | 0.000 | 0.011 | 0.060 |
| *Parabacteroides* genus abundance | 0.001 | 0.003 | 0.001 | 0.015 | 0.002 | 0.002 | 0.087 | 0.048 | 0.043 | 0.096 | 0.001 |
| Time for tumour to treble in volume | 6 | 7 | 10 | 4 | - | 5 | 5 | 9 | 7 | 7 | 14 |
| Time to culling (days) | 14 | 19 | 31 | 10 | 38 | 12 | 10 | 17 | 17 | 19 | 26 |

| Diet | **Insoluble high fibre** | | | | | | | | | | |
|---|---|---|---|---|---|---|---|---|---|---|---|
| Cage | 5 | | | | | | 6 | | | | |
| ID | C5.3 | C5.30 | C5.1 | C5.10 | C5.NM | C5.4 | C6.NM | C6.1 | C6.3 | C6.30 | C6.10 |
| IR/Non-IR | Non-IR | IR | Non-IR | IR | IR | IR | IR | IR | IR | IR | Non-IR |
| *B.acidifaciens* abundance | 0.060 | 0.081 | 0.022 | 0.095 | 0.001 | 0.095 | 0.019 | 0.023 | 0.078 | 0.116 | 0.024 |
| *Parabacteroides* genus abundance | 0.098 | 0.069 | 0.012 | 0.035 | 0.248 | 0.044 | 0.014 | 0.036 | 0.007 | 0.010 | 0.003 |
| Time for tumour to treble in volume | 6 | 8 | 20 | 8 | 6 | 10 | 10 | 10 | 10 | | 11 |
| Time to culling (days) | 17 | 17 | 42 | 19 | 10 | 17 | 19 | 17 | 21 | - | 19 |

| Diet | **Soluble high fibre** | | | | | | | | | | |
|---|---|---|---|---|---|---|---|---|---|---|---|
| Cage | 7 | | | | | | 8 | | | | |
| ID | C7.1 | C7.10 | C7.3 | C7.NM | C7.30 | C7.4 | C8.NM | C8.1 | C8.10 | C8.3 | C8.30 |
| IR/Non-IR | Non-IR | IR | IR | IR | IR | IR | IR | Non-IR | IR | IR | Non-IR |
| *B.acidifaciens* abundance | 0.000 | 0.000 | 0.000 | 0.001 | 0.331 | 0.000 | 0.221 | 0.000 | 0.252 | 0.157 | 0.000 |
| *Parabacteroides* genus abundance | 0.060 | 0.078 | 0.099 | 0.062 | 0.008 | 0.040 | 0.035 | 0.105 | 0.022 | 0.022 | 0.121 |
| Time for tumour to treble in volume | 6 | 10 | 11 | 9 | 12 | 7 | - | 7 | - | - | 12 |
| Time to culling (days) | 10 | 17 | 24 | 17 | 42 | 14 | 38 | 21 | 33 | 42 | 26 |
